# Cookiecutter: a tool for kmer-based read filtering and extraction

**DOI:** 10.1101/024679

**Authors:** Ekaterina Starostina, Gaik Tamazian, Pavel Dobrynin, Stephen O’Brien, Aleksey Komissarov

## Abstract

**Motivation:** Kmer-based analysis is a powerful method used in read error correction and implemented in various genome assembly tools. A number of read processing routines include extracting or removing sequence reads from the results of high-throughput sequencing experiments prior to further analysis. Here we present a new approach to sorting or filtering of raw reads based on a provided list of kmers.

**Results:** We developed Cookiecutter — a computational tool for rapid read extraction or removing according to a provided list of k-mers generated from a FASTA file. Cookiecutter is based on the implementation of the Aho-Corasik algorithm and is useful in routine processing of high-throughput sequencing datasets. Cookiecutter can be used for both removing undesirable reads and read extraction from a user-defined region of interest.

**Availability:** The open-source implementation with user instructions can be obtained from GitHub: https://github.com/ad3002/Cookiecutter.

## 1 Introduction

Next-generation sequencing technologies continuously become cheaper and more available for routine analysis. There are a number of tasks that require extracting or removing specific sequences from raw reads before their assembly. The use of a small fraction of extracted region-specific reads (e.g. from mtDNA or rRNA) can significantly improve the speed and quality of the obtained assembly and its analysis (Hahn *et al*., 2013). On the other hand, trimming of reads is not always a desired strategy and it can be more effective and simpler to remove all reads that contain fragments of technical sequences. Such removing procedure keeps the same read length that is required, for example, for pseudogenome suffix array creation (Kowalski *et al.*, 2015).

Several tools were developed for raw read processing: trimming according to a given library of adapter sequences (Bolger *et al.*, 2014; Martin, 2011), removing sequence contamination according to coverage and alignment identity (Schmieder and Edwards, 2011a), read mapping to reference sequences (Morgan *et al.*, 2009), removing reads according to the sequence entropy measure (Schmieder and Edwards, 2011b) and removing reads according to their copy number (Brown *et al.*, 2012). However, there is a lack of tools for filtering reads according to their similarity to arbitrary sequences. To solve this problem we developed the Cookiecutter tool.

Cookiecutter is a standalone command-line tool package that can be easily integrated into any analysis pipeline. Cookiecutter takes as input one or several FASTQ files or one or several pairs of FASTQ files and requires a file with a list of k-mers for filtering. That list can be either provided by a user or generated by Cookiecutter from a provided FASTA file. Cookiecutter can process both single-end and paired-end reads. For paired-end reads, Cookiecutter maintains read pairs if both reads pass the filtration; if one read of the pair was filtered out, then the other one is output as a single-end read.

## 2 Implementation

### 2.1 Cookiecutter overview

Cookiecutter can perform several kinds of tasks and provides a convenient command-line interface to launch its subprograms.

1. **remove** compares reads to a given list of k-mers and outputs reads without any matches;
2. **rm reads** is an extended version of **remove** that can filter (G)n and (C)n tracks, short reads, reads containing unknown nucleotides and use the DUST filter to detect low-complexity sequences; its output includes both filtered and unfiltered reads;
3. **extract** compares reads to a given list of k-mers and outputs reads that have matches;
4. **separate** outputs both matched and unmatched reads to separate files;
5. **make library** creates a file of k-mers from sequences of a specified FASTA file. Cookiecutter can process multiple input files in parallel mode.

### 2.2 Aho-Corasick string matching algorithm

Cookiecutter is based on the Aho-Corasick string matching algorithm (Aho and Corasick, 1975) implemented in C++ for working with large datasets. The Aho-Corasick string matching algorithm is a dictionary-matching algorithm that locates elements of a finite set of kmers within an input sequence. The main advantage of this algorithm for read filtration or extraction is that it matches all provided kmers simultaneously. Time complexity of the matching procedure in the text of size *m* is *O*(*m* + *k*), where *k* is the number of matches. Since Cookiecutter searches only for the first match and stops the search if a match is found, its time complexity is *O*(*m*), that is, linear in the length of a searched sequence.

### 2.3 Low complexity detection

The symmetric DUST algorithm (Morgulis *et al.*, 2006) was designed to mask low-complexity sequence reads. In Cookiecutter, DUST is employed to filter reads instead of masking nucleotides within them, so the original algorithm was modified to calculate the DUST score for the whole read. Unlike the original implementation that uses triplet counts, the Cookiecutter DUST module allows to specify the size of read subwindows. The rolling hash function was implemented for time-efficient calculation of subwindow counts.

## 3 Common usage scenarios

To estimate Cookiecutter’s running time, we used the Illumina “Platinum genome” dataset (PCR-free *×*30 coverage sequence of the NA12878 human individual, run ERR194147) that is publicly available from the Illumina website http://www.illumina.com/platinumgenomes/. We also used run SRR100173 to demonstrate rRNA contamination removal. All computations were performed on a Dell R815 PowerEdge server with a RAID 6 disk storage that uses Seagate Barracuda 4 Tb hard disks and Raidix software. We found that in all cases the limited bandwidth of the storage system was a bottleneck for fast raw read processing. A Python script for reproducing these results with an arbitrary pair of FASTA files is available in the demo directory of Cookiecutter. Estimated time and bandwidth values for each proposed Cookiecutter usage scenarios are shown in Table 1.

**Table 1:**
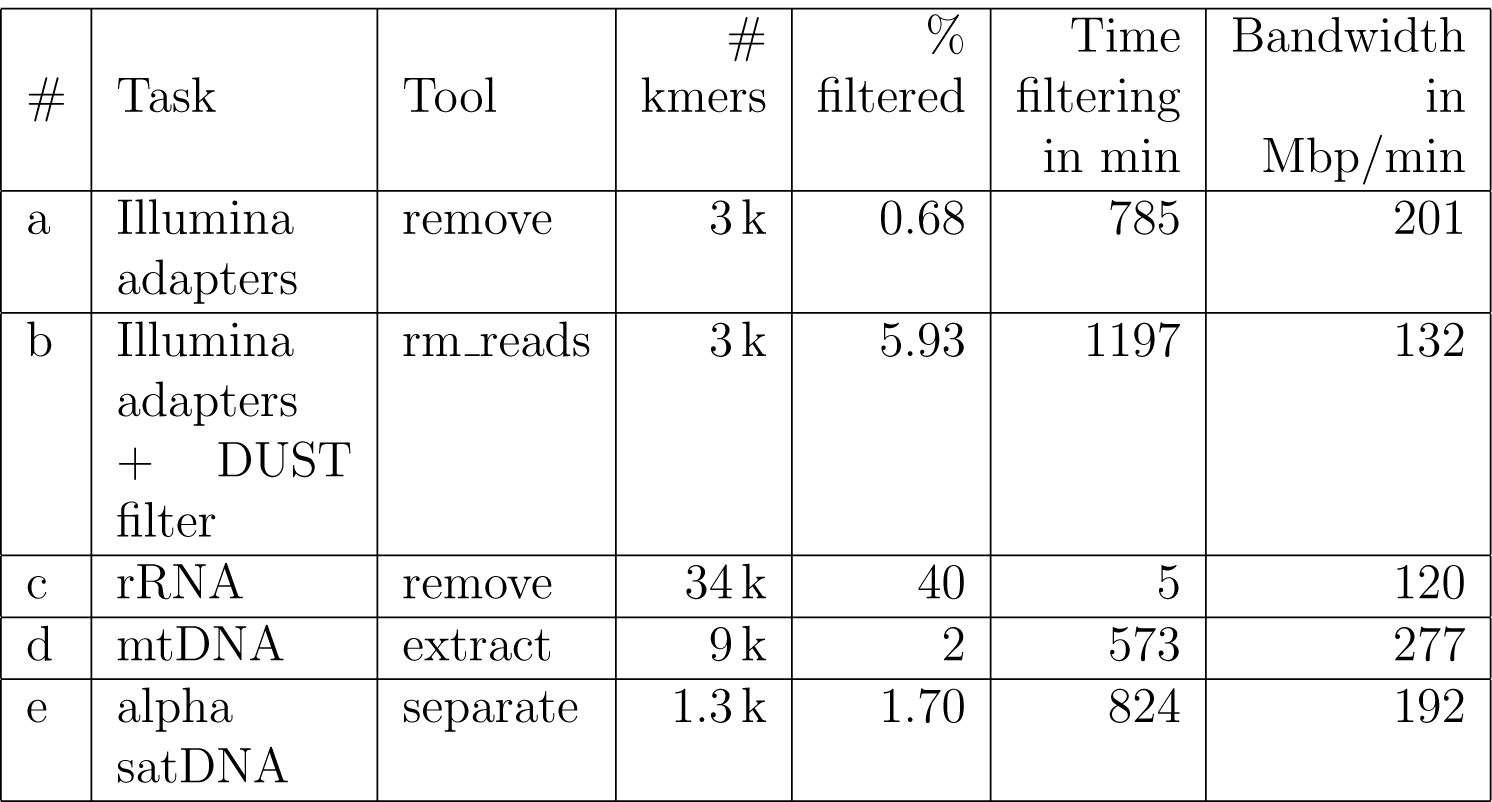
Typical Cookiecutter tasks for NA12878 ×30 Illumina HiSeq dataset. a — Removing Illumina adapters from sequenced library; b — removing Illumina adapters and low-complexity regions from sequenced library; c — removing rRNA reads from transcriptome dataset; d — separating mtDNA reads and whole genome reads from the sequenced library; e — removing alpha satellite DNA reads from the sequenced library.

### 3.1 Removing reads containing technical sequences

The most straightforward application of Cookiecutter is the removal of reads containing any technical sequences during the final preparation of high-quality reads for assembly. These technical sequences can include adapter sequences, Illumina’s indexes and custom PCR primer sequences. By default, Illumina sequencers remove all technical fragments, but they might leave them in a small fraction of reads due to possible errors that could occur during automatic trimming. In such case it is preferable to remove these reads completely. If a dataset contains a large fraction of reads with technical fragments, Cookiecutter can be run after a conventional read trimming procedure to ensure that no technical sequences remain (Table 1a). Additionally, Cookiecutter can remove: reads containing (C)n and (G)n tracks that can frequently be found in Nextra-based datasets; low-complexity reads using the DUST filter and reads containing more than the specified number of unknown nucleotides; reads with length less than a specified length (Table 1b).

### 3.2 Contamination removal

Another Cookiecutter application is the removal of a known contamination. For example, it can remove human DNA contamination from bacterial samples, *Mycoplasma* sp. DNA contamination from a sequenced cell culture or rRNA contamination from transcriptome data. (NCBI, 2015). Cookiecutter can also extract reads that contain target-specific k-mers from a contaminated dataset. For both of these applications a user need to provide a correct k-mer library that contains a non-redundant set of k-mers specific to the contamination and/or target sequences. We used the RNA-Seq dataset SRR100173 contaminated by rRNA as an example of contaminated reads (Table 1c).

### 3.3 Extraction reads from a region of interest

The most prominent application of Cookiecutter is the extraction of reads that contain k-mers speficic to a region of interest (e.g. mitochondrial DNA, a set of genes from a related organism or virus DNA). The extracted reads can be assembled with commonly used assemblers (Table 1d). However, a sequence from a region of interest must be masked for non-specific repeats and low-complexity subsequences to minimize the number of false-positive filtered reads. Iterative usage of Cookiecutter provides the in silico basis for the in vitro prime walking method of extraction of reads flanked by a known region of interest (Chinault and Carbon, 1979).

### 3.4 Smart digital normalization

Cookiecutter can be used for smart digital normalization (Brown *et al.*, 2012) with a given list of high copy number repeats (e.g. centromeric satellite DNA tandem repeats), that might otherwise increase memory and CPU usage during the assembly process (Table 1e).

## 4 Conclusion

We developed an open source software, Cookiecutter, that is useful in routine processing of high-throughput datasets. Cookiecutter can be used for both the removal of undesirable reads and read extraction with a user-defined region of interest. The extraction functionality can be an effective alternative or a complete replacement for read-mapping based pipelines.

